# Dietary sugar and protein exert opposing effects on key larval growth and metabolic regulators, the *Drosophila* insulin-like peptides Dilp2 and Dilp6

**DOI:** 10.1101/2022.12.01.518785

**Authors:** Miyuki Suzawa, W. Kyle McPherson, Elizabeth E. Van Gorder, Shivani Reddy, Dalton L. Hilovsky, Cami N. Keliinui, Leila A. Jamali, Michelle L. Bland

## Abstract

Nutrient intake drives secretion of insulin and insulin-like peptides that stimulate glucose uptake, nutrient storage, protein synthesis and cell growth. The *Drosophila* genome encodes seven insulin-like peptides (Dilps) that bind to a single known insulin receptor to drive growth and nutrient storage. Whether Dilps respond uniformly to changes in dietary nutrients is unknown. Here we characterized the endocrine response to starvation and dietary sugar and protein in mid-third instar *Drosophila* larvae, measuring circulating Dilp2, derived from insulin-producing cells in the brain, and Dilp6, produced by the fat body. Starvation led to a 90% reduction in circulating Dilp2 without affecting circulating Dilp6 levels. Dietary protein, but not sugar, restored hemolymph Dilp2 from starved levels, while elevated and imbalanced ratios of sugar to protein led to modest reductions in circulating Dilp2. In contrast, hemolymph Dilp6 was increased by a sugar-only diet. Surprisingly, dietary protein strongly reduced circulating Dilp6 levels. Dietary sugar drives glycogen and triglyceride storage, and levels of these stored nutrients positively correlate with Dilp6. Protein in the diet promotes whole-animal growth, which correlates strongly with circulating Dilp2. Our data show that Dilp2 and Dilp6 secretion are regulated in opposite ways by distinct dietary nutrients. These findings raise the question of how the single known insulin receptor integrates divergent signals from distinct Dilps to control growth and metabolism.

## INTRODUCTION

The larval stage of the *Drosophila* life cycle is characterized by rapid growth and high levels of nutrient storage (Church and Robertson 1966). These anabolic processes prepare the animal for metamorphosis and adult life (Storelli *et al*. 2019). *Drosophila* insulin-like peptides (Dilps) drive growth and nutrient storage in the larval stage by binding to the single known insulin receptor (InR) in target organs and activating the canonical phosphatidylinositol-3 kinase-Akt signaling pathway (Böhni *et al*. 1999; Verdu *et al*. 1999; Brogiolo *et al*. 2001; Grönke *et al*. 2010). Seven Dilps that bind to and activate InR are encoded in the *Drosophila* genome and expressed in different tissues and at different levels across development. Insulin-producing cells (IPCs) in the fly brain synthesize and release Dilps 2, 3, and 5 into hemolymph as well as at synapses throughout the central brain. Dilp6 is produced and secreted by the fat body (Okamoto *et al*. 2009; Slaidina *et al*. 2009; Suzawa *et al*. 2019). During the larval third instar stage, transcript levels of *Dilp2* decline while *Dilp6* is markedly induced, driven in part by high levels of ecdysone that precede and trigger metamorphosis (Okamoto *et al*. 2009).

In addition to their unique tissue expression patterns and developmental profiles, Dilp2 and Dilp6 exhibit opposite responses to environmental stressors such as starvation and infection. In starved larvae, Dilp2 and Dilp5 are retained in insulin-producing cells, suggestive of decreased circulating levels of these hormones (Géminard *et al*. 2009). In contrast, *Dilp6* transcript levels in larval fat body are induced by starvation, suggestive of increased circulating Dilp6 in starved larvae (Slaidina *et al*. 2009). Activation of the Toll innate immune signaling pathway in the larval fat body strongly suppresses circulating levels of Dilp6, without altering circulating Dilp2 (Suzawa *et al*. 2019).

Whether and how Dilp2 and Dilp6 respond to starvation and specific dietary nutrients and direct responses to nutrient stress is unknown. Here we show that circulating levels of Dilp2 are markedly reduced by starvation, while Dilp6 levels are maintained. We find that Dilp2 and Dilp6 exhibit opposite responses to dietary sugar and protein. Sugar-only diets suppress Dilp2 but lead to elevated levels of Dilp6 in circulation. However, dietary protein promotes secretion of Dilp2 but strongly reduces hemolymph Dilp6 levels. These endocrine changes parallel nutrient storage driven by sugar and whole-animal growth that is promoted by dietary protein. Importantly, our findings also reveal that Dilp2 and Dilp6 are not only regulated by sugar and protein individually, but that each hormone responds to the ratio of these major components in the diet.

## RESULTS

To determine how distinct Dilps respond to nutrient stress, we starved mid-third instar *Drosophila* larvae for increasing periods of time on 0.8% agar and measured hemolymph Dilp2 and Dilp6 as well as transcript levels of *Dilp2* and *Dilp6* in brain and fat body, respectively. In agreement with studies showing that Dilp2 and Dilp5 are retained in insulin-producing cells in starved larvae (Géminard *et al*. 2009), hemolymph Dilp2 was significantly reduced by starvation, with 60% and 84% decreases in circulating levels at 6 and 24 hours of starvation, respectively (Figure 1A). This parallels the nearly complete transit of food through the gut after 6 hours on agar (Figure S1). Transcript levels of *Dilp2* in brain also decreased, but only after 24 hours of starvation (Figure 1B). In contrast with Dilp2, hemolymph levels of Dilp6 were largely unchanged by starvation at all points examined, although we noted a high degree of variability at each time point likely related to differences in larval developmental timing among samples (Figure 1C). As previously reported (Slaidina *et al*. 2009), we find that starvation led to increased *Dilp6* transcript levels in fat body, with increased expression from 6 to 24 hours of starvation (Figure 1D). Starvation rapidly depleted whole-larval glycogen stores, with a significant decrease after just 2 hours of starvation and a 5.4-fold reduction in glycogen after 24 hours of starvation (Figure 1E). In contrast to glycogen, whole-animal triglyceride levels were unchanged over the course of 24 h of starvation on agar (Figure 1F). Together, these data suggest a strong correlation between loss of circulating Dilp2, but not Dilp6, and impaired carbohydrate storage in the response to starvation in *Drosophila* larvae.

**Figure 1.**
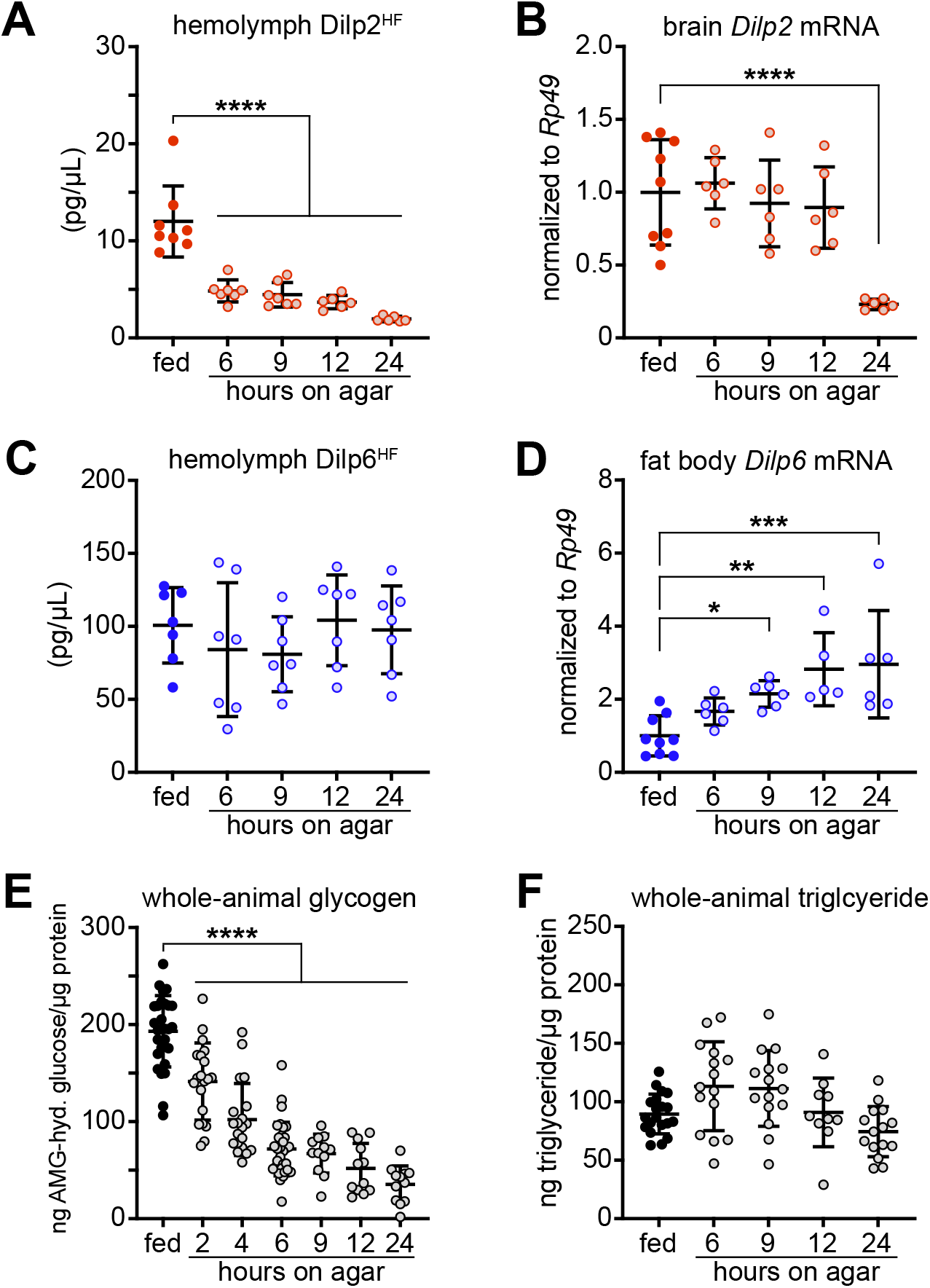
Starvation in third instar *Drosophila* larvae decreases circulating levels of Dilp2 but not Dilp6. Mid-third instar larvae were fed normal food for 24 h or 0.8% agar for indicated times. **A)** Hemolymph Dilp2^HF^ levels in *dilp2*^*1*^, *gdilp2*^*HF*^ larvae, n = 6-8 samples/group. ****p < 0.0001 versus fed. **B)** Whole-brain *Dilp2* transcripts in wild type larvae, normalized to *Rp49*, n = 5-9 samples/group. ****p < 0.0001. **C)** Hemolymph Dilp6^HF^ levels in *dilp6*^*HF*^ larvae, n = 7 samples/group. **D)** Fat body *Dilp6*, measured by RT-qPCR and normalized to *Rp49*, n = 5-9 samples/group. *p = 0.0446, **p = 0.0013, ***p = 0.0003, ****p < 0.0001 versus fed. Hemolymph levels of Dilp2 and Dilp6 were measured in pooled samples collected from 5-6 fed or starved wild type larvae per sample. Transcript levels of *Dilp2* and *Dilp6* were measured in pooled samples from 5-6 brains or 2-3 fat bodies per sample, respectively. **E)** Whole-animal glycogen levels (expressed as amyloglucosidase (AMG)-hydrolyzable glucose), normalized to protein, in fed and starved wild type larvae, n = 12-32 larvae/group. ****p < 0.0001 versus fed. **F)** Whole-animal triglyceride levels, normalized to protein, in fed and starved wild type larvae, n = 10-20 larvae/group. For all panels, larvae were not sorted for sex. Data are presented as means ± SD; p values were determined by one-way ANOVA with Dunnett’s multiple comparison test.

Our data indicate that Dilp2 and Dilp6 respond in markedly different ways to nutrient depletion, but our measurements may be confounded by the known developmental changes in the transcript levels of these hormones. Indeed, the third larval instar in *Drosophila melanogaster* is marked by changes in endocrine factors known to regulate metabolism and growth. For example, measurements of transcript levels in whole larvae show increasing *Dilp6* mRNA levels and declining *Dilp2* and *Dilp5* transcript levels during the third instar stage (Okamoto *et al*. 2009; Slaidina *et al*. 2009; Suzawa *et al*. 2019). These observed hormonal changes and the variability in developmental timing even among larvae hatched within hours of each other prompted us to calibrate time in the third instar using Sgs3-GFP, a fusion of the Salivary gland secretion 3 (Sgs3) and GFP proteins driven under control of the *Sgs3* promoter (Biyasheva *et al*. 2001). As previously reported, expression of Sgs3-GFP is first detectable at ∼100 hours after egg lay (h AEL), midway through the third instar. Changes in pattern and intensity of GFP expression over the next 20-24 hours allow staging of larvae at ∼112 and ∼120 h AEL (Figure S2A). We measured Dilp transcripts and hemolymph levels in third instar larvae at three distinct Sgs3-GFP stages (A: ∼100 h AEL, B: ∼112 h AEL, C: ∼120 h AEL). In agreement with published findings (Slaidina *et al*. 2009), whole-brain *Dilp2* transcripts decreased during the third instar, and we found similar stage-dependent reductions in hemolymph Dilp2, with 60% lower hemolymph Dilp2 levels at 120 h AEL compared with 100 h AEL (Figure S2B and S2C). Whole-brain mRNA levels of *Dilp5* paralleled *Dilp2*, but *Dilp3* transcript levels were steady throughout the second half of the third instar (Figure S2D and S2E). In contrast, we find stepwise increases in both fat body *Dilp6* transcripts and circulating Dilp6 levels as larvae progress through the third instar, with a 4.5-fold increase in hemolymph Dilp6 from 100 to 120 h AEL (Figure S2F and S2G). The changes in circulating Dilp levels over the larval third instar underscore the need for synchronizing larvae to achieve reliable measurements of hormones such as Dilp2 and Dilp6.

We examined responses of Dilp2, Dilp6, circulating sugar and stored nutrients to starvation stress in larvae staged using Sgs3-GFP. We fed larvae normal fly food or 0.8% agar for 24 hours beginning in the early-to mid-third instar, and after 24 hours, we scored Sgs3-GFP expression in fed and starved larvae, selecting those with expression intermediate to stage A and stage B (∼106-108 h AEL). This ensured that while a group of third instar larvae with varying developmental time entered the experiment, the animals used for measurements had reached the same developmental point after 24 hours of feeding or starvation. Additionally, this approach ensured that larvae used for hormone, metabolite and growth measurements were subject to starvation beginning at ∼84 h AEL, after critical weight had been achieved (BEADLE *et al*. 1938; Mirth and Riddiford 2007). We find that starvation reduced hemolymph Dilp2 levels by 77-80% in male and female larvae, respectively (Figure 2A). Starvation led to non-significant, 26-40% reductions in circulating Dilp6 in both sexes (Figure 2B). Hemolymph trehalose levels were reduced by 20-24% in starved larvae compared with fed animals (Figure 2C), with no change in hemolymph glucose (Figure 2D). Starvation significantly depleted glycogen stores by 2-fold in male larvae and 2.3-fold in females (Figure 2E) but had no significant effect on triglyceride storage in males or females (Figure 2F). Starvation of larvae beginning from ∼84 h AEL to the end of the larval stage significantly reduced whole-animal growth by 14% in males and 18% in females compared with fed animals. Transferring staged larvae that had been starved for 24 hours back to food for the remainder of the third instar increased growth by 3.5% in males and 8.6% in females versus larvae that remained on agar (Figure 2G and 2H).

**Figure 2.**
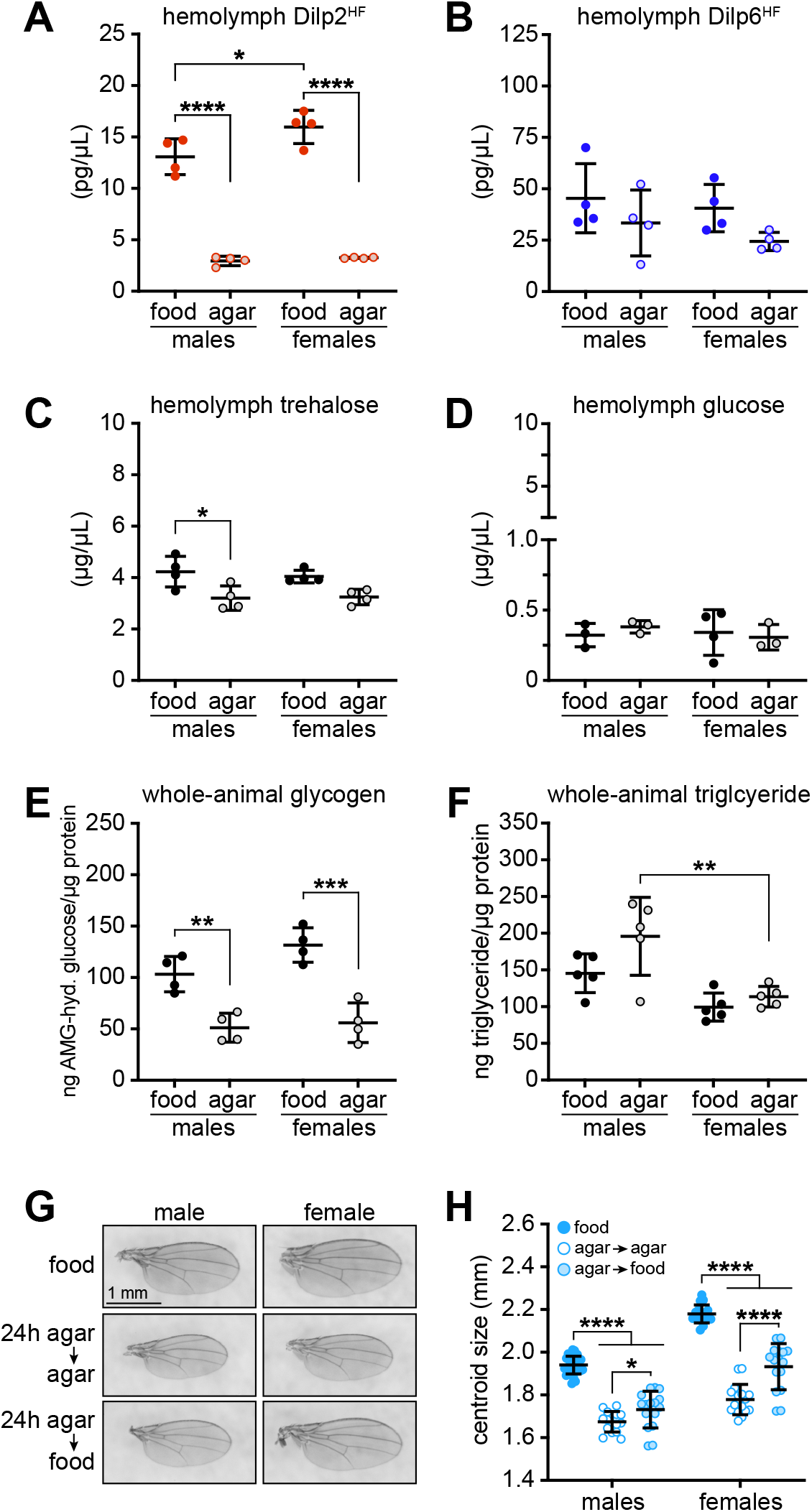
Endocrine, metabolic, and growth effects of starvation are due to nutrient depletion and not alterations in developmental timing. Early-to mid-third instar larvae carrying *Sgs3-GFP* were fed normal food or starved on 0.8% agar for 24 h, and those with salivary gland GFP fluorescence indicating progression to ∼108 h AEL were selected for further analysis. **A)** Hemolymph Dilp2^HF^ in male and female *dilp2*^*1*^, *gdilp2*^*HF*^, *Sgs3-GFP* larvae, n = 4 samples/group. *p = 0.0236, ****p < 0.0001. **B)** Hemolymph Dilp6^HF^ in male and female *dilp6*^*HF*^; *Sgs3-GFP* larvae, n = 4 samples/group. **C)** Hemolymph trehalose and **D)** hemolymph glucose in male and female larvae, n = 4 samples/group. *p = 0.0235. **E)** Whole-animal glycogen levels, normalized to protein, in male and female larvae, n = 4 larvae/group. **p = 0.0045, ***p = 0.0002. **F)** Whole-animal triglyceride levels, normalized to protein, in male and female larvae, n = 5 larvae/group. **p = 0.0044. **G)** Representative images of wings from adult male and female flies that were reared as larvae on normal food (top) or were starved on agar for 24 h, sorted for salivary gland fluorescence indicating progression to ∼100 h AEL, and then transferred to vials containing agar (center) or normal food (bottom) for the remainder of the larval and pupal stages. Scale bar, 1 mm. **H)** Adult wing centroid size, n = 16-30 wings/group. *p = 0.0178, ****p < 0.0001. Data are presented as means ± SD; p values were determined by one-way ANOVA with Tukey’s multiple comparison test.

Given the distinct responses of Dilp2 and Dilp6 to starvation, we asked whether these hormones are also differentially regulated by dietary nutrients. In the wild and in the laboratory, the two major components of the *Drosophila* diet are plant sugars and yeast. Dietary yeast is the main protein source for fruit fly larvae, and it also supplies cholesterol and phospholipids (Carvalho *et al*. 2012). We used a nutritional geometry approach to determine how dietary sugar and protein regulate Dilps, nutrient storage and growth (Lee *et al*. 2008; Skorupa *et al*. 2008; Post and Tatar 2016). We fed larvae normal fly food or diets containing sucrose or yeast extract, with each nutrient added at 0%, 1% or 10% (w/v) in 0.8% agar and in varying combinations for 24 hours from the mid-third instar until ∼108 h AEL, determined using Sgs3-GFP reporter fluorescence. Larvae that reached this stage after 24 hours on experimental diets were used for further analyses.

Insulin-producing cells in the larval brain release Dilp2 and Dilp5 in response to dietary protein (Géminard *et al*. 2009). In agreement with previous reports, we find that addition of yeast extract to plain agar media substantially and dose-dependently increases Dilp2 levels in hemolymph compared with starved larvae. In larvae fed diets with 1% or 10% sucrose but no added yeast extract, circulating Dilp2 levels remained low and indistinguishable from the starved condition. At the highest level of dietary yeast tested (10%), addition of sucrose at 10% reduced hemolymph Dilp2 by 27-32% compared with diets that contained 0% or 1% sucrose (Figure 3A). Dilp2 levels responded similarly to dietary nutrients in male and female larvae, but females exhibited higher circulating Dilp2 levels than males in fed and 10% yeast with 0% or 1% sucrose conditions (p ≤ 0.05, Student’s unpaired t tests).

**Figure 3.**
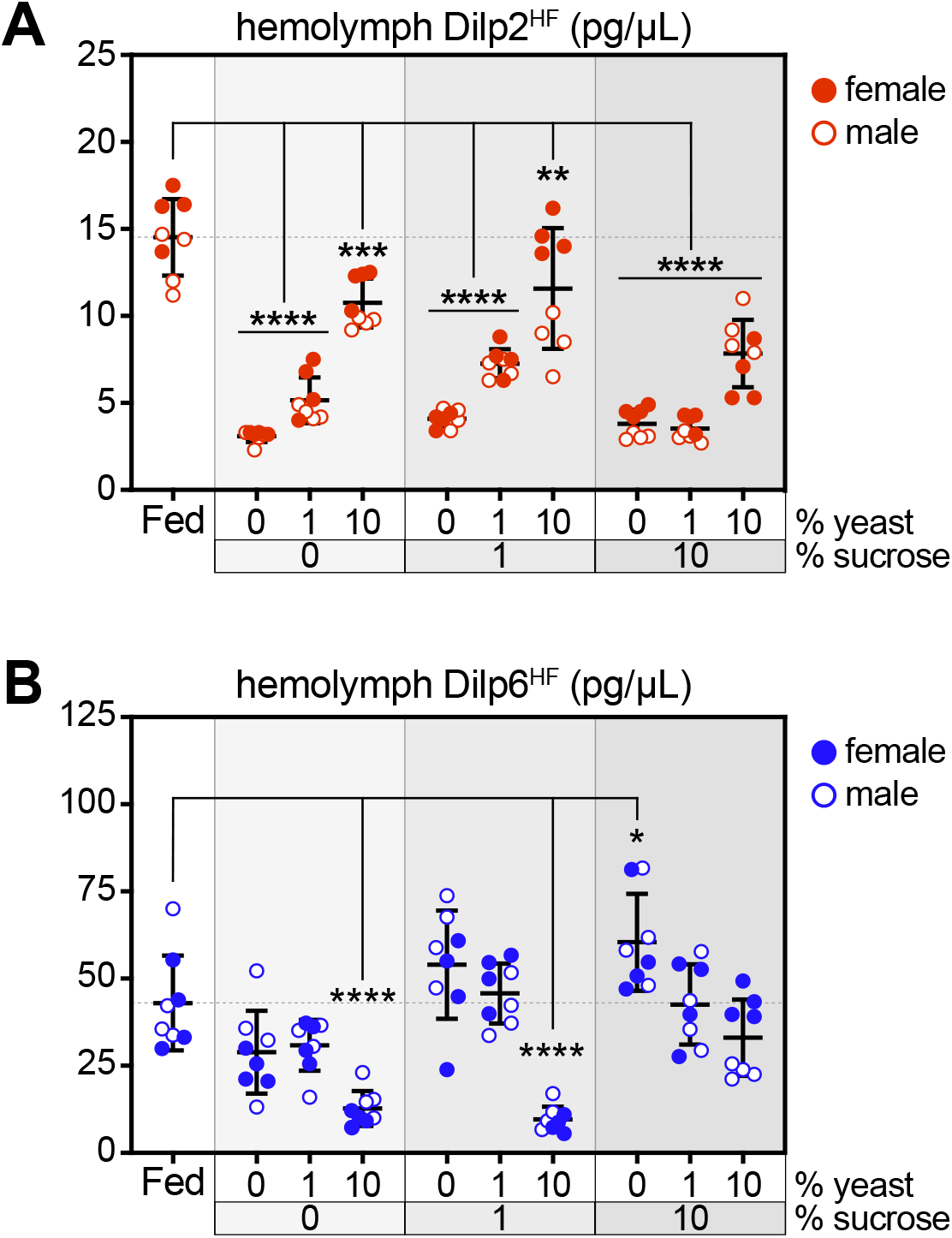
Dilp2 and Dilp6 exhibit opposing responses to dietary nutrients. Early-to mid-third instar larvae carrying *Sgs3-GFP* were fed normal food, starved on 0.8% agar, or fed indicated concentrations of sucrose and/or yeast extract in 0.8% agar for 24 h, and those with salivary gland GFP fluorescence indicating progression to ∼108 h AEL were selected for further analysis. **A)** Hemolymph Dilp2^HF^ in *dilp2*^*1*^, *gdilp2*^*HF*^, *Sgs3-GFP* male and female larvae. **B)** Hemolymph Dilp6^HF^ in *dilp6*^*HF*^; *Sgs3-GFP* male and female larvae. n = 4 samples/sex/group, with 8 samples total/group. *p = 0.0145, **p = 0.0042, ***p = 0.001, ****p < 0.0001 versus fed. Fed and 0% sucrose, 0% yeast data in panels A and B are identical to food and agar data in Figure 2A and 2B, respectively. Gray shading in graphs indicates sucrose dose; male and female samples are indicated by open and filled symbols, respectively. Data are presented as means ± SD; p values were determined by one-way ANOVA with Dunnett’s multiple comparison test.

Dietary nutrients exerted opposite effects on Dilp6 compared with Dilp2. Larvae fed a diet containing 10% yeast extract in agar exhibited a 71% decrease in hemolymph Dilp6 compared with fed animals. In contrast, addition of sucrose to agar media increased circulating Dilp6 in a dose-dependent manner, leading to 1.25- and 1.4-fold increases in Dilp6 levels relative to larvae consuming normal fly food. Addition of 10% yeast extract to diets containing 10% or 1% sucrose reduced hemolymph Dilp6 to levels at or below those observed in larvae fed normal fly food (Figure 3B). Dilp6 levels responded similarly to dietary nutrients in male and female larvae, but females exhibited higher circulating Dilp6 levels than males in the 10% yeast, 10% sucrose condition (p = 0.0002, Student’s unpaired t test). In summary, our data show that in *Drosophila* larvae, Dilp2 is suppressed by starvation and restored by dietary protein but not sugar. Dilp6 is unaffected by starvation, down-regulated in response to dietary protein, and induced by dietary sugar.

The insulin signaling pathway is a key regulator of carbohydrate, lipid, and protein metabolism across the animal kingdom. We examined levels of key metabolites in larvae raised for 24 hours during the mid-third instar on normal fly food, starvation media, or agar with varying levels of sucrose and yeast extract, using Sgs3-GFP to identify animals that had reached the same developmental stage for measurements. Hemolymph levels of trehalose, the major circulating sugar in *Drosophila*, were highly responsive to dietary conditions (Figure 4A). Larvae fed food without sucrose exhibited 17-32% decreases in hemolymph trehalose compared with animals fed normal fly food. In contrast, experimental diets with sucrose only led to dose-dependent, 18-22% increases in circulating trehalose. Introducing yeast extract to diets with 1% or 10% sucrose reduced circulating trehalose, similar to the pattern observed with hemolymph Dilp6. Glucose also circulates in *Drosophila* larvae, but at ten-fold lower levels compared with trehalose. In larvae fed experimental diets for 24 h in the mid-third instar, we observed no changes in circulating glucose with the exception of a small increase in animals fed 1% yeast extract in agar (Figure 4B).

**Figure 4.**
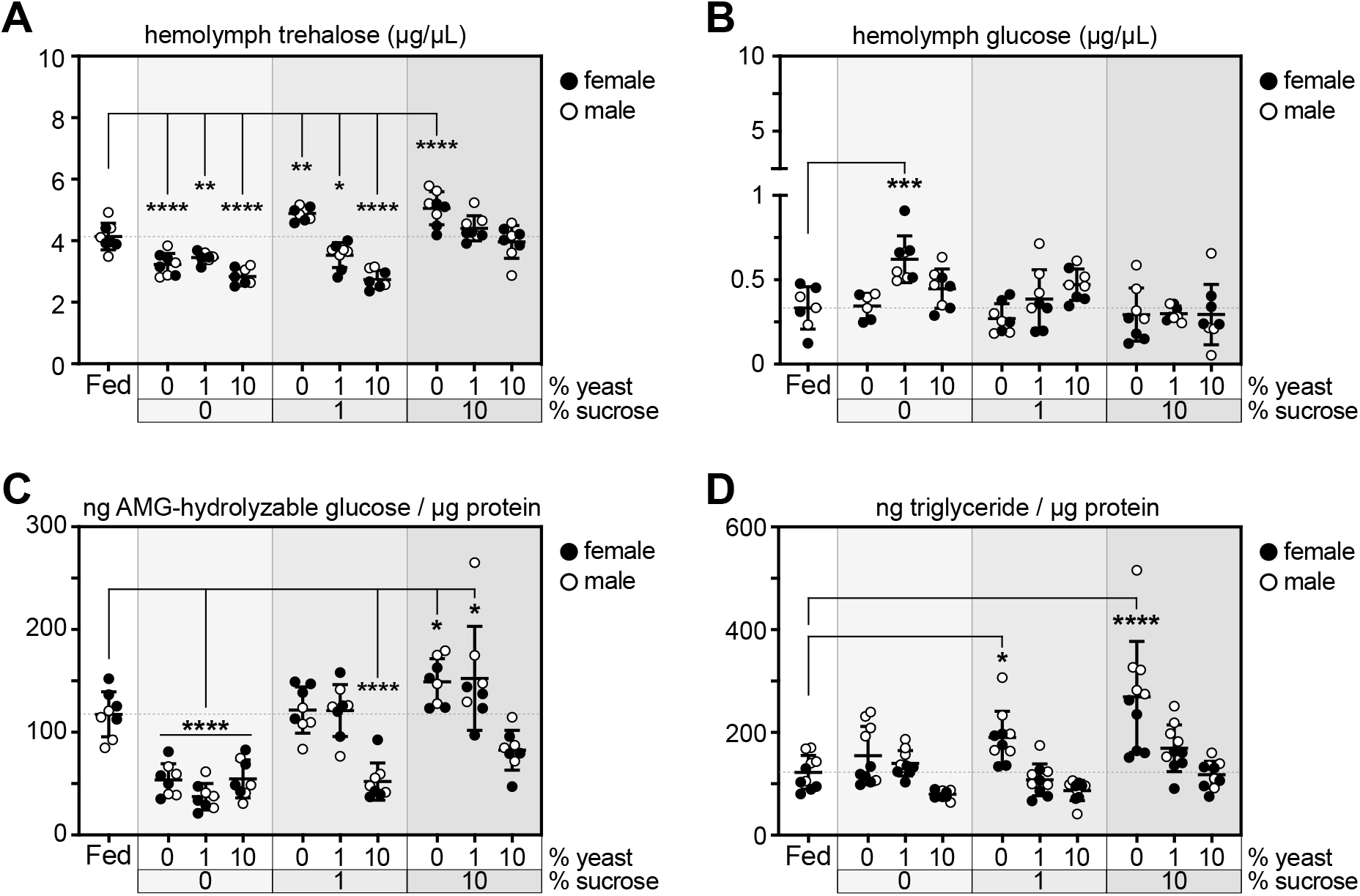
Dietary sugar drives nutrient storage. Early-to mid-third instar larvae carrying *Sgs3-GFP* were fed normal food, starved on 0.8% agar, or fed indicated concentrations of sucrose and/or yeast extract in 0.8% agar for 24 h, and those with salivary gland GFP fluorescence indicating progression to ∼108 h AEL were selected for further analysis. **A)** Hemolymph trehalose and **B)** hemolymph glucose levels in male and female larvae, measured in samples pooled from 6-8 larvae, n = 4 samples/sex/group, with 8 total samples/group. *p = 0.0161, **p ≤ 0.0044, ***p = 0.0004, ****p < 0.0001 versus fed. **C)** Whole-animal glycogen levels, normalized to total protein, in male and female larvae, n = 4 larvae/sex/group, with 8 total larvae/group. *p ≤ 0.0435 and ****p < 0.0001 versus fed. **D)** Whole-animal triglyceride levels, normalized to total protein, in male and female larvae, n = 5 larvae/sex/group, with 10 total larvae/group. *p = 0.0168 and ****p < 0.0001 versus fed. Fed and 0% sucrose, 0% yeast data in panels A-D are identical to food and agar data in Figure 2C-2F, respectively. Gray shading in graphs indicates sucrose dose; male and female samples are indicated by open and filled symbols, respectively. Data are presented as means ± SD; p values were determined by one-way ANOVA with Dunnett’s multiple comparison test.

Dietary sugar is stored as glycogen in the larval fat body and somatic musculature and as triglyceride in fat body. Like circulating trehalose, glycogen storage was driven by dietary sugar. In mid-third instar larvae fed diets with yeast extract only for 24 hours, glycogen levels, normalized to whole-body protein levels, were reduced 2.1-to 3.1-fold versus larvae fed normal fly food. Diets with 1% sucrose completely rescued glycogen storage to fed levels, and diets with 10% sucrose elevated glycogen levels by 27-30% relative to levels in animals fed normal fly food (Figure 4C). We noted an apparent suppressive effect of high levels of dietary protein (10% yeast) on glycogen storage in larvae fed 1% and 10% sucrose. However, we note that whole-animal protein levels are increased in animals fed 10% yeast, thereby altering calculation of glycogen stores when normalized to protein (Figure S3A). Dietary sugar also drove fat storage in a dose-dependent manner. Larvae fed diets with increased sucrose for 24 hours exhibited 1.6- and 2.2-fold increases in triglyceride levels, normalized to whole-animal protein, relative to larvae fed normal food. We noted no major effects of the other experimental diets on triglyceride to protein ratios (Figure 4D). However, as was the case for glycogen measurements, whole-animal protein levels were highly responsive to dietary yeast, affecting calculations of triglycerides measured per lysate volume (Figure S3B).

Insulin-like peptides are major regulators of whole-animal growth in *Drosophila* (Grönke *et al*. 2010). We evaluated whole-animal growth in larvae fed normal fly food, starved on agar, or fed experimental diets with varying amounts of sucrose and/or yeast extract from the mid-third instar (∼84 h AEL) through the pupal stage or for 24 hours in the mid-third instar stage (∼84 to ∼108 h AEL). In each case, after 24 hours on a given diet, larvae were sorted for salivary gland Sgs3-GFP fluorescence indicating a developmental stage equivalent to ∼108 h AEL, after which they were returned to the experimental diet or to normal fly food. Following eclosion, wings were dissected from adult flies and centroid size was measured as a proxy for whole-animal growth (Figure 5A). Starvation of larvae from the mid-third instar through the pupal stage strongly inhibited wing growth by 14-18% compared with flies fed normal food. However, addition of 10% yeast extract alone to agar media rescued wing size to 95-97% of that in flies fed normal food during the third instar. Diets containing 10% sucrose alone only partially rescued animal growth compared with starvation media, leading to wings that were 10-13% smaller than those from animals fed normal food. The experimental diet containing 10% sucrose and 10% yeast extract led to an almost complete rescue of growth compared with the fed condition (Figure 5B and Figure S4).

**Figure 5.**
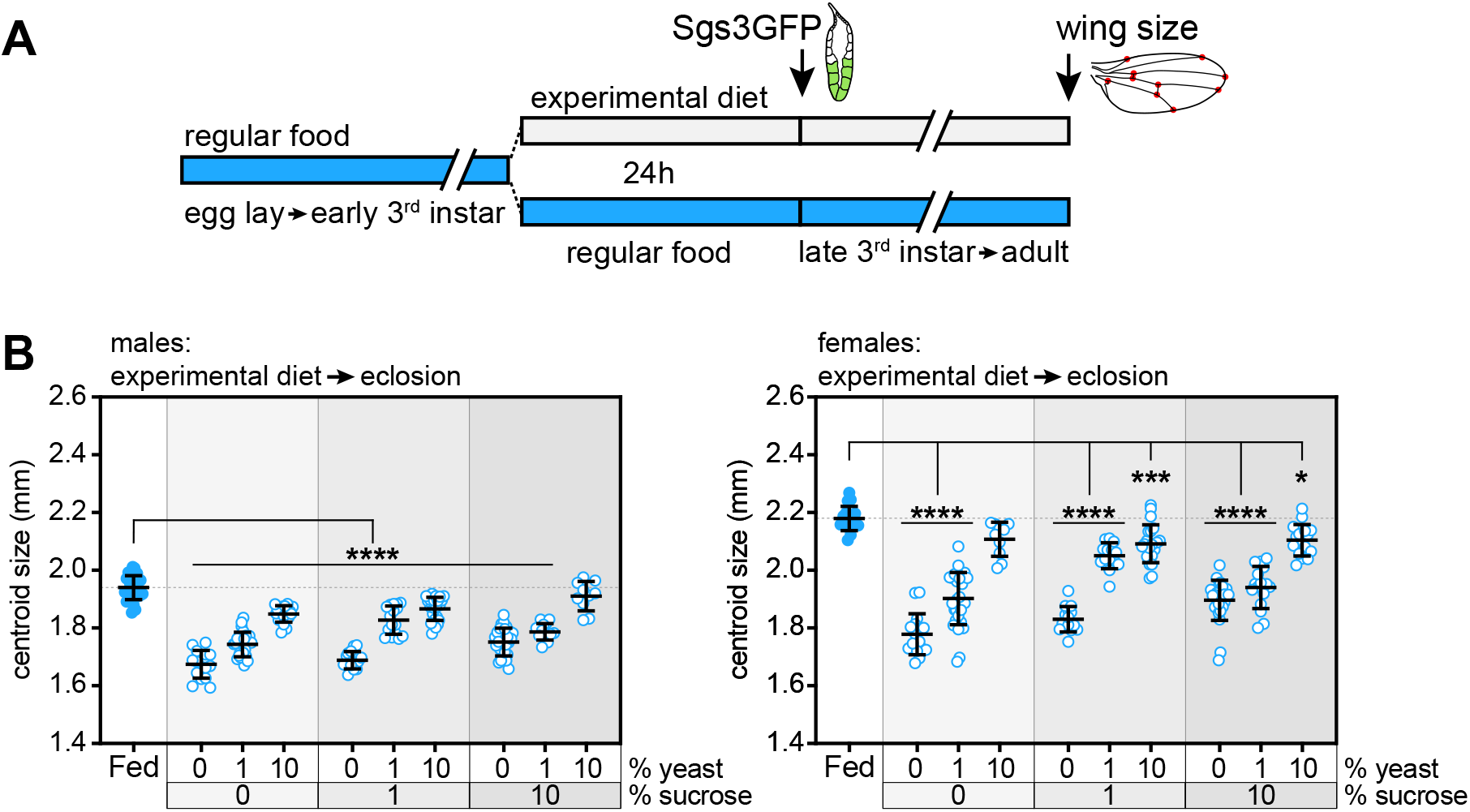
Dietary protein drives whole-animal growth. **A)** Early-to mid-third instar larvae carrying *Sgs3-GFP* were fed normal food, starved on 0.8% agar, or fed indicated concentrations of sucrose and/or yeast extract in 0.8% agar for 24 h, and those with salivary gland GFP fluorescence indicating progression to ∼108 h AEL were transferred to vials containing the same experimental diet for the remainder of the larval and pupal stages. Centroid size was measured in wings dissected from adult flies 2-3 days after eclosion. **B)** Adult wing centroid size in male (left) and female (right) flies reared on normal food (fed) or experimental diets from mid-third instar to eclosion, n = 10-31 wings/group. *p = 0.0110, ***p = 0.0006, ****p < 0.0001 versus fed. Fed and 0% sucrose, 0% yeast data in panels B are identical to food and agar data in Figure 2H. Gray shading in graphs indicates sucrose dose. Fed data are identical for graphs in B as the same animals served as controls in both experiments. Data are presented as means ± SD; p values were determined by one-way ANOVA with Dunnett’s multiple comparison test.

## DISCUSSION

Here we used a nutritional geometry approach to discover how the major components of the Drosophila diet, sugar and yeast, regulate circulating levels of two insulin receptor ligands, Dilp2 and Dilp6. Based on our findings, Dilp2 is sensitive to starvation and likely regulated by dietary yeast but not sucrose. Dilp6 is insensitive to starvation and, unexpectedly, it is positively regulated by dietary sucrose and negatively regulated by yeast intake. Our data also show that animal growth, like Dilp2 but not Dilp6, is highly responsive to dietary protein. Furthermore, we find that storage of triglycerides and especially glycogen strongly correlates with dietary sucrose, much like Dilp6.

Our data show that hemolymph levels of Dilp2 and Dilp6 change in opposite directions not only in response to dietary nutrients but also over the course of the larval third instar. Indeed, this critical stage of *Drosophila* development is marked not only by endocrine changes but also by rapid growth and nutrient storage. We used a fluorescent reporter of salivary gland developmental progression to precisely stage third instar larvae during the last half of the larval third instar. This allowed us to compare animals that had achieved an equivalent developmental stage after 24 hours on various diets in our nutritional geometry approach, enabling reliable measurements of Dilp2, Dilp6, nutrient storage and growth.

Our results highlight important roles for dietary sucrose in *Drosophila* larvae. Sucrose is frequently added to minimal media for either fruit fly larvae or cultured larval organs for starvation (Britton and Edgar 1998; Géminard *et al*. 2009). We find that plain agar and agar that contains sucrose yield clearly distinct phenotypes with regard to hormone levels, nutrient storage and growth. Starvation via agar feeding suppresses Dilp2 but not Dilp6 and reduces glycogen storage. In contrast, media containing only 1% sucrose leads to elevated Dilp6 levels and drives glycogen accumulation compared with plain agar lacking any nutrients.

Our data and data from other labs raise a key question: how does the single insulin receptor (InR) encoded in the *Drosophila* genome interpret the information carried by distinct Dilps? One possibility is that InR is simply responding to the total level of Dilps in circulation. This seems unlikely given the large differences in Dilp2 and Dilp6 levels in larvae fed 10% yeast or 10% sucrose and the effects of these diets on growth. Another possibility is that Dilp2 and Dilp6 exhibit different affinities for insulin-binding peptides that restrict or promote their access to InR perhaps in a tissue-specific manner. Whether accessory proteins in Dilp target tissues might bias InR sensitivity to one Dilp over another is unknown. This scenario would most closely resemble the condition in mammals, where insulin and IGF1 bind to distinct receptors on their target organs.

Our findings show that Dilps exhibit divergent responses to the most critical components of an animal’s environment: availability of key dietary nutrients. Dilp2 and Dilp6 are highly sensitive to distinct dietary nutrients with large changes in the level of each hormone accompanying the presence or absence of sucrose or yeast. The exact information carried by each hormone and the targets of its action remain to be discovered.

## FIGURE LEGENDS

**Supplementary Figure 1.**
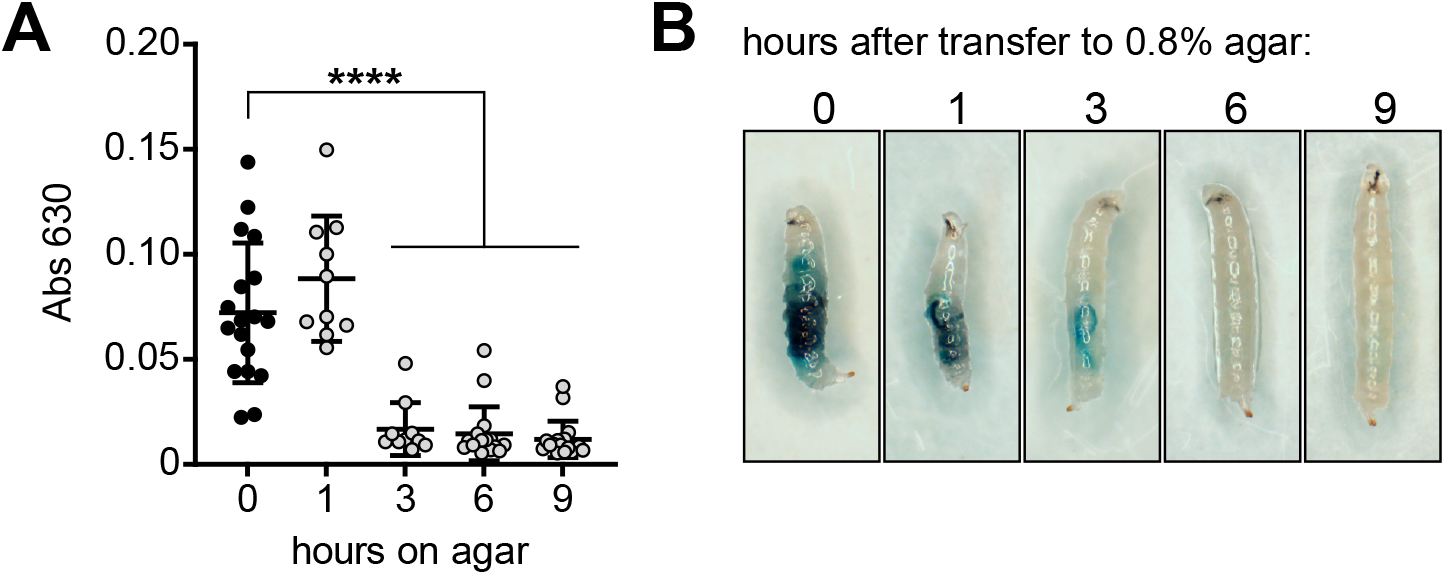
Progression of ingested food through the gut in starved larvae. Mid-third instar larvae reared on FD&C No. 1 blue-dyed food were transferred to agar for indicated periods of time to monitor progression of ingested food through the gut. **A)** Quantitation of blue food in larval lysates by absorbance readings at 630 nm, n = 10-18 larvae/group. ****p < 0.0001 versus 0 h. **B)** Representative images of larvae at each time point. Larvae were not sorted for sex. Data are presented as means ± SD; p values were determined by one-way ANOVA with Dunnett’s multiple comparison test.

**Supplementary Figure 2.**
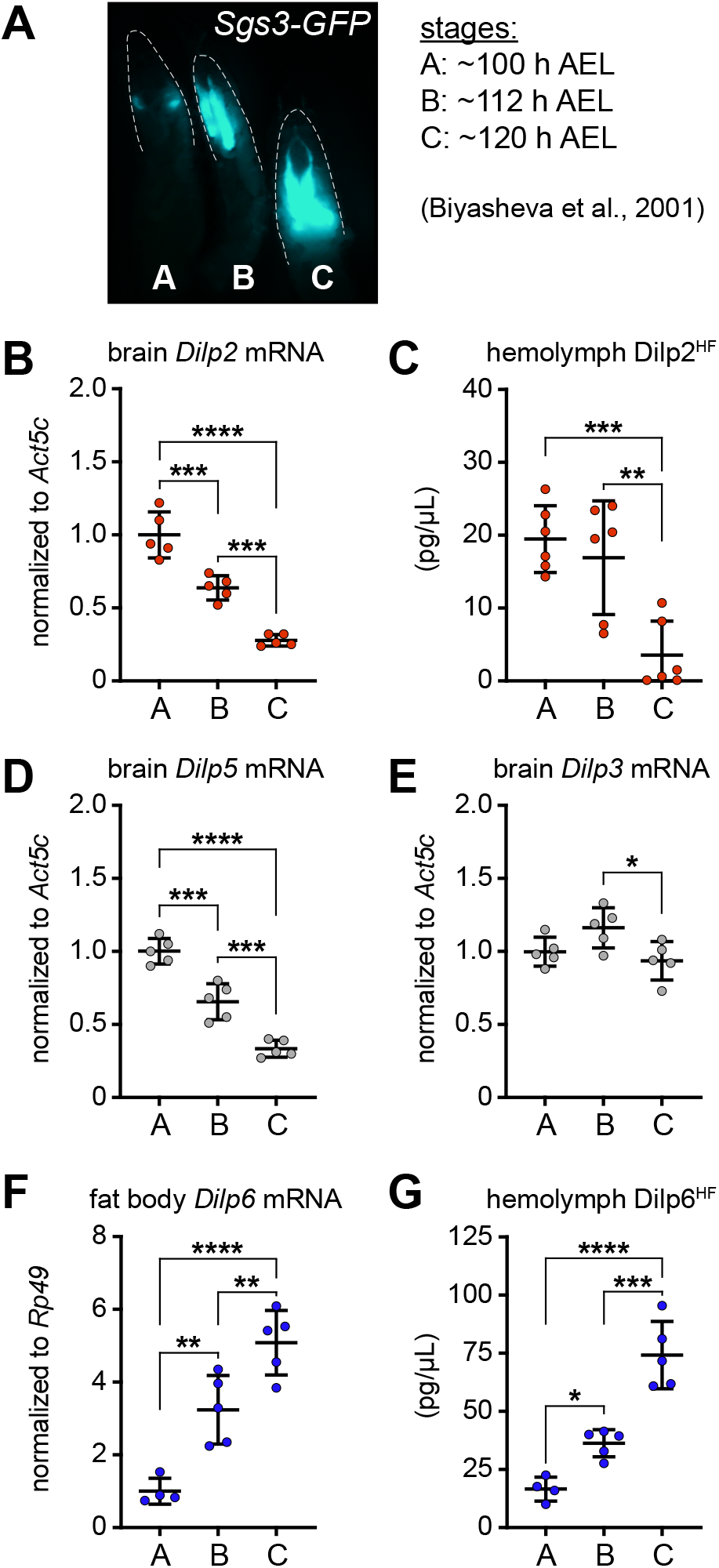
Sgs3-GFP allows staging of the last 24 hours of the third instar and calibration of Dilp levels over developmental time. Fluorescent micrograph of *Sgs3-GFP* transgenic third instar larvae. Three easily distinguishable salivary gland GFP expression patterns allow staging from ∼100-120 h AEL (at 25°C) (Biyasheva *et al*. 2001). Stage A: partial expression at ∼100 h AEL. Stage B: full expression at ∼112 h AEL. Stage C: full expression plus GFP in salivary gland lumen at ∼120 h AEL. For panels B-F, larvae carrying *Sgs3-GFP* were collected at stages A, B and C as described above. **B)** Whole-brain *Dilp2* transcripts in wild type larvae, normalized to *Act5c*, n = 5 samples/group. **C)** Hemolymph Dilp2^HF^ levels in *dilp2*^*1*^, *gdilp2*^*HF*^, *Sgs3-GFP*, n = 6 samples/group. **D)** Whole-brain *Dilp3* and **E)** *Dilp5* transcripts in wild type larvae, normalized to *Act5c*, n = 5 samples/group. **F)** Fat body *Dilp6* transcripts in wild type larvae, normalized to *Rp49*, n = 4-5 samples/group. **G)** Hemolymph Dilp6^HF^ levels in *dilp6*^*HF*^; *Sgs3-GFP* larvae, n = 4-5 samples/group. *p ≤ 0.0339, **p ≤ 0.01, ***p ≤ 0.0008, ****p < 0.0001, as indicated. For all panels, larvae were not sorted for sex. Data are presented as means ± SD; p values were determined by one-way ANOVA with Tukey’s multiple comparison test.

**Supplementary Figure 3.**
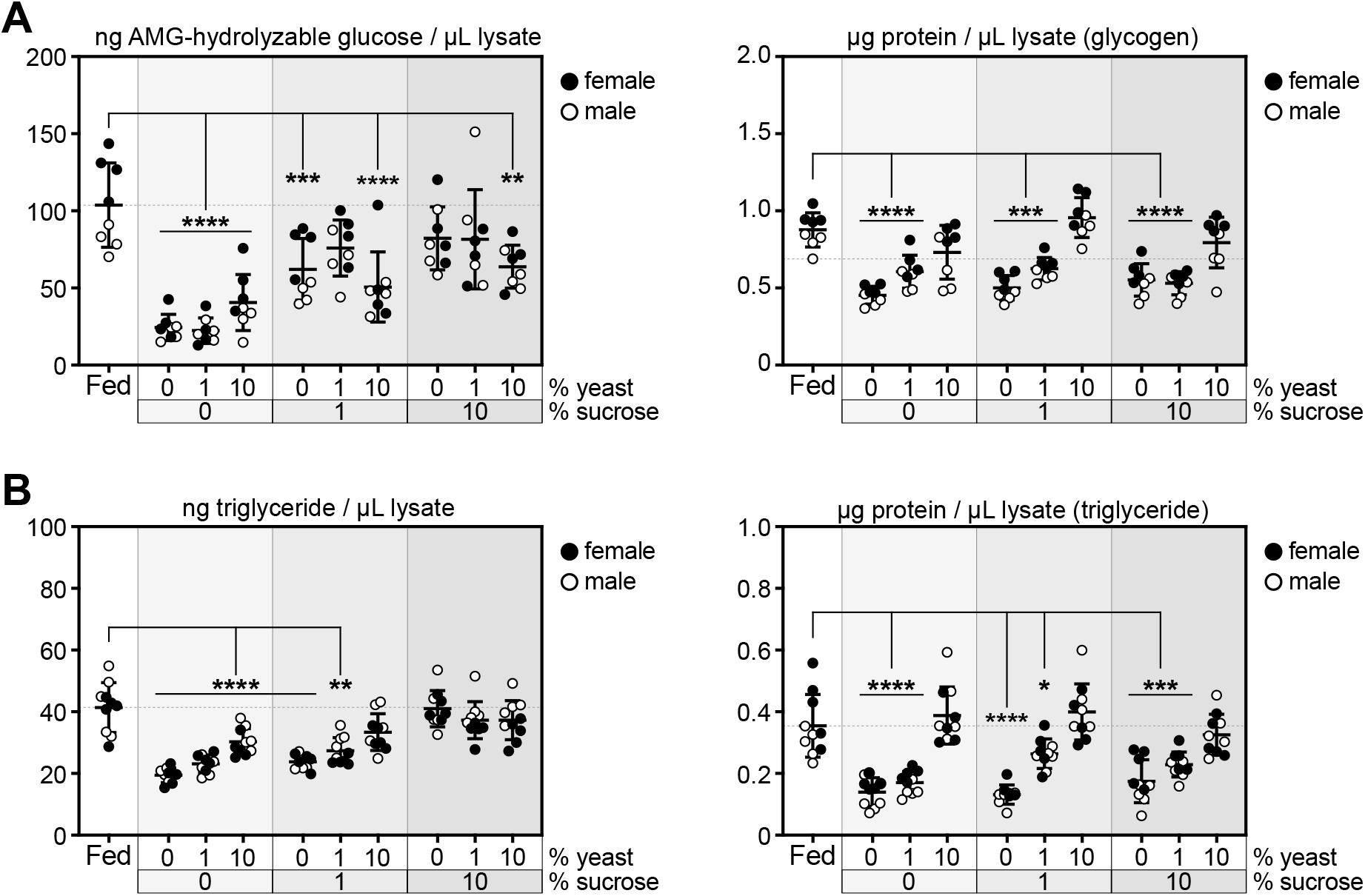
Dietary sugar drives nutrient storage and dietary yeast drives whole-animal protein levels. Early-to mid-third instar larvae carrying *Sgs3-GFP* were fed normal food, starved on 0.8% agar, or fed indicated concentrations of sucrose and/or yeast extract in 0.8% agar for 24 h, and those with salivary gland GFP fluorescence indicating progression to ∼108 h AEL were selected for further analysis. **A)** Left: whole-animal glycogen levels, normalized to lysate volume, in male and female larvae. Right: whole-animal protein levels, normalized to lysate volume, in samples used for glycogen measurement. n = 4 larvae/sex/group with 8 total larvae/group. **p = 0.0015, ***p ≤ 0.0009, ****p < 0.0001 versus fed. **B)** Left: whole-animal triglyceride levels, normalized to lysate volume, in male and female larvae. Right: whole-animal protein levels, normalized to lysate volume, in samples used for triglyceride measurement. n = 5 larvae/sex/group with 10 total larvae/group. *p = 0.0228, **p = 0.0066, ***p = 0.0005, ****p < 0.0001 versus fed. Gray shading in graphs indicates sucrose dose; male and female samples are indicated by open and filled symbols, respectively. Data are presented as means ± SD; p values were determined by one-way ANOVA with Dunnett’s multiple comparison test.

**Supplementary Figure 4.**
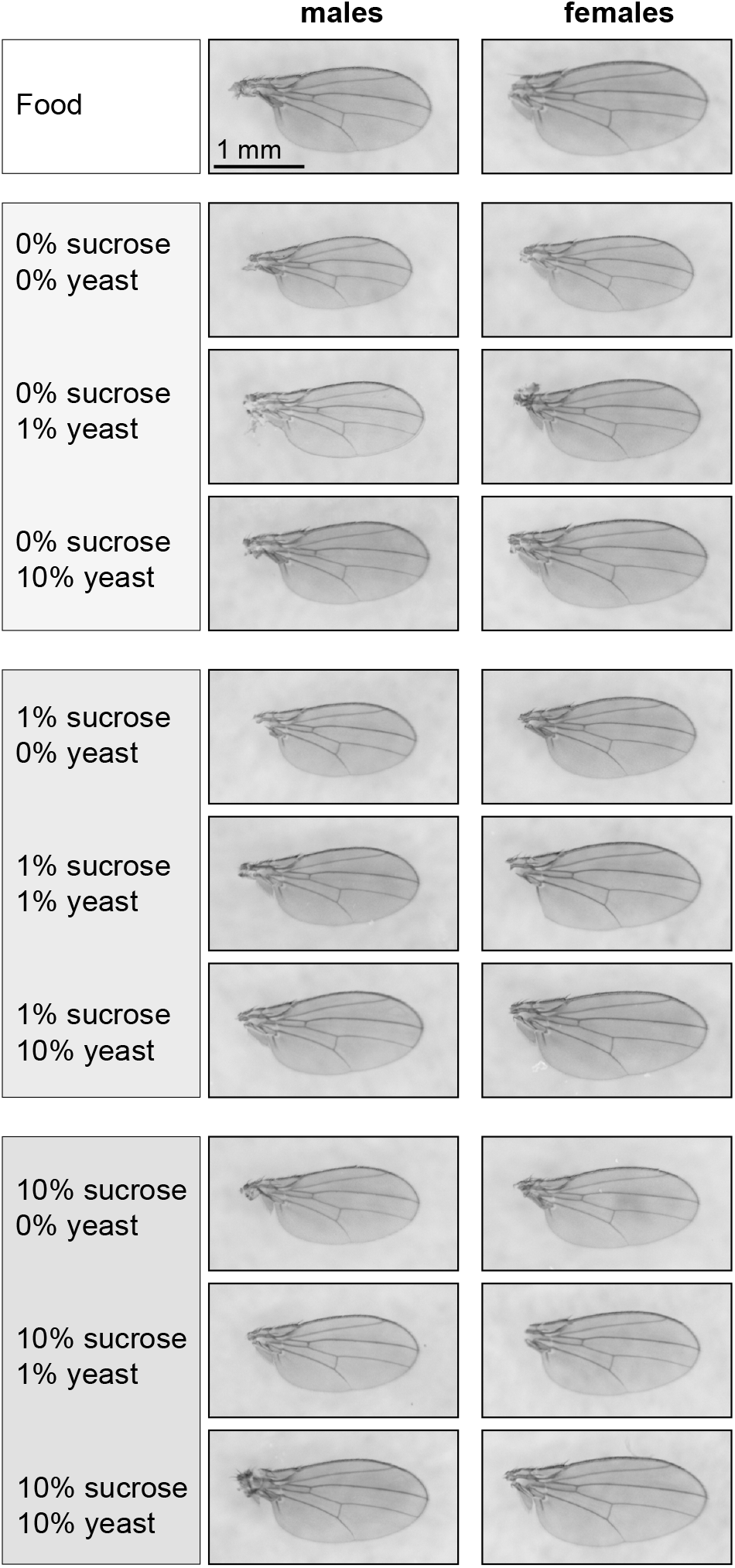
Representative images of wings from flies grown on experimental diets. Wings were dissected from 2-3 day old adult male and female flies that were grown on each of ten diets for 24 hours from the early-to mid-third instar stage, selected for salivary gland GFP fluorescence indicating progression to ∼108 h AEL, and transferred to vials containing the same diet (experimental diet) or normal fly food (experimental diet→ food) for the remainder of the larval and pupal stages. Note that the “food” wing images are identical for each dietary regimen as the same animals are used as controls for each experiment. Representative images were chosen on the basis of centroid values closest to the experimental mean of each group. Scale bar, 1 mm.

## METHODS

### *Drosophila* stocks and husbandry

Flies were raised on food containing 7.8% molasses, 2.4% yeast, 4.6% cornmeal, 0.3% propionic acid, and 0.1% methylparaben (Archon Scientific, Durham, NC). Experiments were performed using larvae that were in the early-to mid-third instar stage (∼80-90 h after egg lay) at the beginning of each feeding regimen. For nutrient addback experiments, sucrose (Fisher BP220) or yeast extract (Fisher BP9727) were added at 1% or 10% concentrations to 0.8% agar (Fisher BP1423). For starvation and experimental diet feeding experiments, mid-third instar larvae were floated off food with 2M NaCl, rinsed in PBS and transferred to food, agar, or agar containing nutrients as indicated. Larvae were fed diets for 24 hours at a density of 12-24 larvae/well in 12-well plates containing 1.5 mL food/well and capped with foam stoppers. For staging using Sgs3-GFP, larvae were gently removed from experimental diets to PBS, and salivary gland GFP fluorescence was evaluated. Larvae at ∼108 h AEL (refer to Figure S2) were selected, sorted for sex, and collected for further analyses. The following stocks were obtained from the Bloomington *Drosophila* Stock Center (Bloomington, IN): isogenic *w*^*1118*^ for experiments in wild type larvae (#5905) and Sgs3-GFP (#5885). Other flies used were: *dilp2*^*1*^, *gDilp2*^*HF*^ (Park *et al*. 2014) and *dilp6*^*HF*^ (Suzawa *et al*. 2019).

### Measurement of hemolymph Dilp levels by ELISA

Hemolymph Dilp levels were measured in pooled samples collected on ice from 5-6 larvae. 1 µL of pooled hemolymph was used for Dilp2^HF^ and Dilp6^HF^ measurement by ELISA using previously established protocols (Park *et al*. 2014; Suzawa *et al*. 2019). Briefly, for ELISA measurements, Immuno Clear plates (Thermo Fisher Scientific) were coated overnight at 4 °C with anti-FLAG antibody (Sigma-Aldrich, F1804) diluted to 5 µg/mL in 0.2 M sodium carbonate/bicarbonate buffer (pH 9.4). Plates were washed with PBS containing 0.2% Tween 20 (PBS-T) and blocked in sterile-filtered 2% bovine serum albumin (BSA) in PBS for 3 h at room temperature. Following washes with PBS-T, FLAG(GS)HA peptide standard (DYKDDDDKGGGGSYPYDVPDYA-amide, 2412 daltons, used at 0.5 – 50 pg/µL) or hemolymph (diluted 1:100 in ice-cold PBS) was added to each well. Anti-HA-Peroxidase 3F10 antibody (Sigma-Aldrich 12013819001), diluted to 15 ng/mL in PBS with 1% Triton X-100, was added to each well for a final volume of 50 µL/well. Plates were incubated in humidified chambers overnight at 4 ºC. After washes with PBS-T, 1-Step Ultra TMB ELISA Substrate (Thermo Fisher Scientific) was added to each well. Plates were incubated for 15 minutes at room temperature, and reactions were stopped by adding 2 M sulfuric acid. Absorbance was measured at 450 nm using an Infinite 200 PRO plate reader (Tecan). To convert molar concentrations to pg/µL hemolymph, we used molecular weights of 7828.86 daltons 10303.62 daltons for mature Dilp2^HF^ and Dilp6^HF^ proteins, respectively.

### Measurement of hemolymph metabolites

Hemolymph was collected on ice from mid-third instar larvae (hemolymph from 5-8 larvae pooled per sample). Endogenous trehalase was destroyed by heating hemolymph diluted in PBS at 70°C for 20 min. The sample was split in half, 1 mU trehalase (Sigma-Aldrich, T8778) was added to one tube, and both were incubated at 37°C for 2 h. Glucose was measured in both samples as described below for glycogen measurements. Trehalose was calculated by subtracting glucose values in trehalase-free samples from glucose values in trehalase-treated samples, and then dividing by two as trehalose is a dimer of glucose.

### Measurement of glycogen and triglyceride levels

For glycogen measurements, larvae were homogenized in ice-cold 0.1 M NaOH using a Kontes pestle, heated at 70°C for 20 min and cleared by centrifugation at 4°C. Samples were incubated for 1h at 37°C with 0.2 M sodium acetate, pH 4.8, with or without amyloglucosidase (5 mg/mL, Sigma-Aldrich 10115). Samples were then incubated for 15 min at room temperature with assay buffer containing glucose oxidase (0.2 U/mL, Sigma-Aldrich G7141), horseradish peroxidase (0.3 U/mL, Sigma-Aldrich P8250) and Amplex UltraRed (25 µM, Invitrogen A36006). Fluorescence in glucose standards (Fisher CAS 50-99-7) and experimental samples was measured (excitation/emission maxima = 535/587 nm) with a Tecan Infinite 200 PRO plate reader. Glycogen levels were calculated by subtracting free glucose (without amyloglucosidase added) from the amount of amyloglucosidase-hydrolyzed glucose and then normalized to protein levels (BCA Protein assay, Pierce). For triglyceride measurements, frozen larvae were sonicated in ice-cold buffer containing 140 mM NaCl, 50 mM Tris-HCl, pH 7.4, 0.1% Triton X-100 with protease inhibitors (Roche). Following clearing by centrifugation at 4°C, glycerol and triglyceride levels in each sample were measured using a Serum Triglyceride Determination Kit (Sigma-Aldrich, components F6428, T2449, and G7793) and normalized to protein levels (BCA Protein assay, Pierce).

### Wing centroid size

Wings were dissected from two-to three-day old adult male and female flies and mounted in Gary’s Magic Mount (1.6 g/mL Canada balsam in methyl salicylate). Images of wings were collected with an Olympus MVX10 microscope and DP73 camera using CellSens software (Olympus). FIJI was used to measure x- and y-coordinates of ten landmarks at the intersections of wing veins with each other or the wing margin. Centroid size was calculated as the square root of the sum of squared distances from each landmark to the wing centroid (Klingenberg and Zaklan 2000).

### Quantitative RT-PCR

Total RNA was extracted from larval fat bodies (2-5 pooled/sample) or larval brains (5 pooled/sample) using the Direct-zol RNA MicroPrep kit (Zymo Research). DNAse-treated total RNA (1 µg) was used to generate cDNA using a High-Capacity cDNA Reverse Transcription kit (Thermo Fisher Scientific). Gene expression was measured using gene-specific primers (listed below). Quantitative PCR reactions were performed on 10-20 ng cDNA using SYBR Select Master Mix (Thermo Fisher Scientific) with a Bio-Rad CFX Connect Real-Time PCR Detection System. Relative amounts of transcripts were calculated using the comparative Ct method with *Rp49* or *Act5c* as a reference gene (Schmittgen and Livak 2008).

### Primers for RT-qPCR

Primers are listed 5’→ 3’.

Dilp6-F ATATGCGTAAGCGGAACGGT

Dilp6-R GCAAGAGCTCCCTGTAGGTG

Dilp2-F ACGAGGTGCTGAGTATGGTGTGCG

Dilp2-R CACTTCGCAGCGGTTCCGATATCG

Dilp3-F GTCCAGGCCACCATGAAGTT

Dilp3-R AAGTTCACGGGGTCCAAAGTT

Dilp5-F TGGACATGCTGAGGGTTGC

Dilp5-R CCGCCAAGTGGTCCTCATAAT

Rp49-F CGCTTCAAGGGACAGTATCTG

Rp49-R AAACGCGGTTCTGCATGA

Act5c-F GCACCGTCGACCATGAAGAT

Act5c-R CTCGTCGTACTCCTGCTTGG

### Quantitation and statistical analysis

Statistical parameters including exact sample sizes (with n referring to biological replicates), data plotted (mean ± SD), p values, and statistical tests used are reported in Figure Legends. Statistical analyses were performed using Graphpad Prism 9. Data were analyzed by Student’s unpaired t test or by one-way ANOVA with the Tukey or Dunnett multiple comparisons test.

## ACKNOWLEDGEMENTS

We thank Claire Dalby (University of Virginia) for help with experiments. We thank Seung Kim (Stanford University) for *dilp2*^*1*^, *gd2*^*HF*^ flies and the Bloomington *Drosophila* Stock Center for other flies used in this study. This work was supported by NIH Grant R01DK123433 to M.L.B. and NIH Pharmacological Sciences Training Grant T32 GM007055 to E.E.V.

## Notes

### Competing Interest Statement

The authors have declared no competing interest.

